# Temporally varying isotopic niche overlap of the invasive ctenophore *Mnemiopsis leidyi* with other zooplanktivores in the western Dutch Wadden Sea

**DOI:** 10.1101/402602

**Authors:** Lodewijk van Walraven, Wouter van Looijengoed, A. Sarina Jung, Victor T. Langenberg, Henk W. van der Veer

## Abstract

The invasive ctenophore *Mnemiopsis leidyi* can be a major predator of zooplankton in areas where it has been introduced. In this study, the possible competition of *M. leidyi* with native macroplankton and nekton in the western Dutch Wadden Sea was investigated in March–August, 2011 by determining and comparing isotopic niches of zooplanktivores. Stable carbon and nitrogen isotope signatures were determined from tissue samples of fish, scyphozoa, hydromedusa, ctenophores, crustaceans and cephalopods. *δ*^15^N of *M. leidyi* was positively related to ctenophore size, suggesting that small ctenophores occupied a lower trophic level than large ones. A cluster analysis showed that in the spring and early summer period, when *M. leidyi* densities are low, average *δ*^13^C and *δ*^15^N ratios of the invasive *M. leidyi* were similar to those of most other gelatinous zooplankton and pelagic fish species sampled. At the beginning of the bloom period in August there was no overlap in isotopic niche of *M. leidyi* with that of any other pelagic zooplanktivore. During this month the population consisted mainly of larvae and juveniles with lower *δ*^15^N ratios. At present, *M. leidyi* appears not to be a significant competitor for other gelatinous zooplankton and fish species because the period of high diet overlap with other consumers was also the period in which *M. leidyi* was least abundant.

## Introduction

*Mnemiopsis leidyi* is an opportunistic planktonic predator of western Atlantic coastal waters which feeds on a wide range of different zooplankton prey such as copepods, copepodites and nauplii, bivalve veligers, barnacle nauplii and cyprids [1,2], fish larvae [3] and eggs [4].

For several decades now, *M. leidyi* has been observed outside of its native range [reviewed in 5]. The first invasion of *M. leidyi* was in the Black Sea, where it was found in 1982. After 1989 density and biomass of the species reached very high levels following recruitment failure in the dominant zooplanktivorous fish species, the anchovy *Engraulis encrasicolus* [6] due to overfishing. A lack of competition, eutrophication and climate induced enhanced carrying capacity led to a competitive advantage of *M. leidyi* over pelagic fish [7]. Fisheries in the Black Sea region suffered substantial economic losses due to the collapse of pelagic stocks [8].

Recently, *M. leidyi* has also been reported from many different areas in western Europe: Sweden [9], Germany [10,11], Denmark [12], Poland [13], Norway [14] and the Dutch coast [15,16] including the western Dutch Wadden Sea where it now occurs year-round in significant numbers, especially in summer and autumn [17].

Previous studies estimated the grazing pressure of gelatinous zooplankton on mesozooplankton prey in the Wadden Sea to be low [18–20]. The introduction of *M. leidyi* warrants a closer evaluation of the role of gelatinous zooplankton as predators of mesozooplankton in the area. In the German part of the Wadden Sea *M. leidyi* shows a high overlap in diet with the zooplanktivorous fish *Clupea harengus* [21] and based on gut content analysis of Wadden Sea fish species, several other species are known to have similar diets to *C. harengus* [22], warranting a closer look at possible competition between *M. leidyi* and other zooplanktivores.

Comparing diets of different animals using gut content analysis can be difficult. Especially jellyfish and other gelatinous zooplankton often have fast digestion rates [23,24], and digestion time varies for different prey species [25]. For instance, in *M. leidyi* digestion times ranged from 0.4 hours for tintinnid ciliates to 4.8 hours for *Acartia tonsa* copepods [1].

In addition to gut content analysis, Stable Isotope Analysis (SIA) has been used for the analysis of diet of gelatinous zooplankton in a wide range of ecosystems [26–29] as well as freshwater and marine fish [26,30–32]. Several studies that compared jellyfish stable isotope ratios with those of pelagic fish showed that the diet of the two groups can overlap [21,26] or reveal fishes as predator of jellyfish [33].

The goal of this study is test the hypothesis that the diet of the invasive ctenophore *Mnemiopsis leidyi* in the western Wadden Sea is similar to that of native gelatinous zooplankton- and fish species by comparing nitrogen and carbon stable isotope signatures of the different species and their overlap with stable isotope signatures of *M. leidyi.* The following research questions are discussed:

- What is the isotopic niche of gelatinous zooplankton and pelagic fish in the Dutch Wadden Sea and does this change over time?
- What is the isotopic niche of *M. leidyi* in the Dutch Wadden Sea and does this change with size?
- Which gelatinous zooplankton species have an overlapping isotopic niche with the invasive ctenophore *Mnemiopsis leidyi*?

Trophic fractionation of *δ*^13^C (Δ*δ*^13^C) and *δ*^15^N (Δ*δ*^15^N) has been shown to be different for different temperatures [34] and diet sources [35]. In experiments with fish [34] as well as jellyfish [36] Δ*δ*^15^N was found to differ from the mean value of 3.4 ‰ used most often for fish [32,37]. As trophic fractionation is unknown for several possible trophic relationships in our study, we did not convert *δ*^15^N to Relative Trophic Position (RTP).

## Materials and methods

### Field sampling

Several types of sampling gears were used to collect samples of gelatinous zooplankton as well as fish in the western Wadden Sea area. Along a transect in the Marsdiep area oblique hauls of 10 minutes were made using an Isaacs- Kidd midwater trawl with a trawling speed of 2 knots. The net had a mouth opening of 4 m^2^ and a mesh size of 5 mm in the back part.

At fixed stations in tidal gullies of the Balgzand intertidal additional samples were taken every one or two weeks using plankton nets made of polyamide plankton gauze (Monodur 2000, 2 mm mesh size) with an opening of 0.7 m^2^, a length of 5 m, a porosity of 0.59 and a total surface area of 12 m^2^. Oblique hauls were made with the ship at anchor in the tidal current. On board catches were sorted and all animals present, or a subsample, were measured to the nearest mm according to Table 1.

**Table 1.** Measurements used for different groups. All measurements were in mm

| group | measurement |
| --- | --- |
| Fish | total length |
| Scyphozoa | bell diameter |
| Athecate hydromedusae | bell height |
| Thecate hydromedusae | bell diameter |
| Beroid and cydippid ctenophores | polar length |
| Lobate ctenophores | oral-aboral length |
| Other invertebrates | total length |

Seasonal patterns, densities and species composition of gelatinous zooplankton in the catches in 2011 as well as 1981 - 1983 and 2009, 2010 and 2012 are available and discussed in work [38].

### Stable isotope analysis

On each survey at least 5 individuals of each species were collected if possible for stable isotope analyses of *δ*^15^N and *δ*^13^C.

Bell tissue was used for scyphomedusae as *δ*^13^C and *δ*^15^N of the bell appears to be representative for that of the whole body in *Aurelia aurita* (d’Ambra et al. 2014). For ctenophores, except for smaller sized *Mnemiopsis leidyi* (<15 mm), tissue around the stomach and statocyst was removed to prevent contamination. *M. leidyi* smaller than 15 mm were placed into filtered seawater for 2–4 hours for digestion of possible food and the whole individual was used. Back muscle tissue was used for fish [39]. For small fish, larvae and some species such as the pipefish *Syngnathus rostellatus* excision of the back muscle tissue was not possible and a section of tail without gonads, stomach and intestines was taken.

In addition, two plankton fractions were collected: seawater was collected using a bucket and filtered over a 80 µm sieve after which the filtrate was filtered through a GF/F filter.

All samples were stored in glass vials at −20 °C until freeze drying. Samples were freeze dried for at least 24 hours until constant weight.

At least 1 mg (fish and non-gelatinous plankton) or 10 mg (gelatinous zooplankton) was put in 9×5 mm tin cups. Freeze-dried and encapsulated samples were analysed for *δ*^13^C and *δ*^15^N SI ratios using a Thermo Scientific Delta V Advantage Isotope Ratio Mass Spectrometer equipped with a Flash 2000 Organic Element Analyser at the Royal Netherlands Institute for Sea Research, Texel, Netherlands.

l-Glutamic acid (*δ*^15^N −4.36 ‰, *δ*^13^C −26.2 5‰, 40.81 % C, 9.52 % N) and

Acetanilide (*δ*^15^N 1,3 ‰ *δ*^13^C −26,1 ‰, 71.08 % C, 10.36 % N) were used as reference material for quantification of C and N ratios, respectively [40,41]. The ^13^C composition was expressed relative to the level in Peedee-Belemnite limestone:

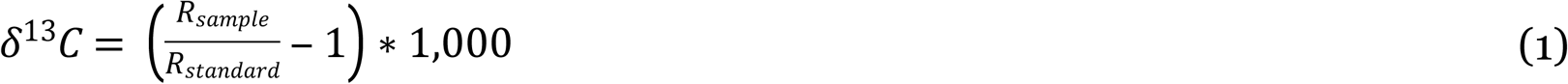

With R being the ratio ^13^C:^12^C. The ^15^N composition was expressed relative to the level in atmospheric N_2_ using the same formula with R being the ratio ^15^N:^14^N.

Samples were analysedin duplicate unless insufficientmaterialwasavailable. Samples from individuals with a standard deviation larger than 1 ‰ for *δ*^13^C (24 samples) and larger than 2.5 ‰ for *δ*^15^N (3 samples) were excluded from the analysis.

*δ*^13^C values were corrected for lipid content as lipid content can influence *δ*^13^C values in aquatic animals, especially at higher lipid contents [42]. The carbon-to nitrogenratio (C:N) by mass of the sample as determined during analysis for *δ*^13^C and *δ*^15^N was used to apply the correction described in [42]:

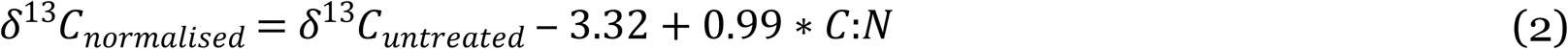

## Data analysis

The primary consumer used as baseline was a filter-feeding bivalve, as used in similar studies [21,32]. Blue mussels *Mytilus edulis*, were collected from the tidal flats of the Mokbaai intertidal area. Additionally on 5-7-2018 *Ensis leei* bivalves were collected at several locations with varying distance to the main freshwater input source, the sluices near the town of Den Oever to investigate difference in *δ*^15^N ratios due to the input of anthropogenic nitrogen.

Stable isotope ratios of consumers were compared in two ways. Average values were compared by performing a cluster analysis, and within- and between- species variation was investigated using the Standard Ellipse Area method as described below.

A cluster analysis was performed on all consumer isotope data collected in 2011 to investigate general patterns and similarity in *δ*^15^N and *δ*^13^C values between species, which has been used in various marine systems before [43–45]. This analysis was similar to the one used in Nagata et al. [45]. A matrix of Euclidean Metric Distances (EMD) was created from mean *δ*^15^N and *δ*^13^C values of each species with the main clusters identified at an EMD of < 2.5.

Variation in stable isotopic niche of consumers was investigated by calculating the Standard Ellipse Area corrected for small sample sizes (SEA_*c*_) from *δ*^15^N and *δ*^13^C measurements of individuals, both over the whole sampled period and per month. This is a bi-variate measure of variation similar to the standard deviation [46].

Possible diet overlap between *Mnemiopsis leidyi* and other consumers was investigated by calculating the percentage of overlap of the area of the SEA_*c*_ ellipse of the various consumers with that of *M. leidyi*. The estimation of SEA_*c*_ and overlaps were performed using the SIBER routines [46] found in the R package SIAR, version 4.2 [47].

To investigate whether the *δ*^15^N ratio of *M. leidyi* differed between ctenophore size, season or both, a series of linear regression models were hypothesised that described the relationship between ctenophore size (oral-aboral length in mm) and *δ*^15^N. In the full model M1, *δ*^15^N was related to ctenophore size and the intercept as well as the slope of this relationship differ per month:

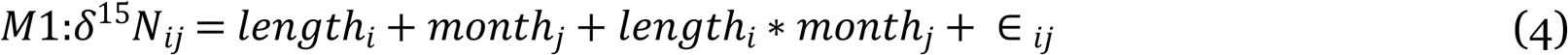

In model M2 *δ*^15^N was related to ctenophore size with a different intercept but similar slope for each month:

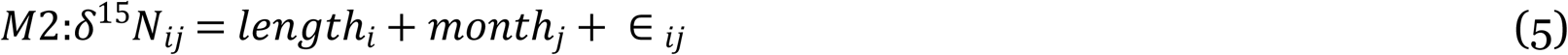

In model M3 *δ*^15^N was related to ctenophore size only:

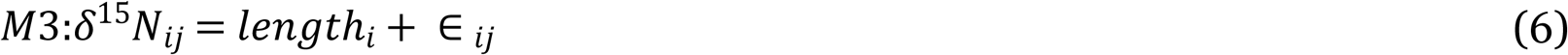

In model M3 *δ*^15^N was not related to ctenophore size but differs per month:

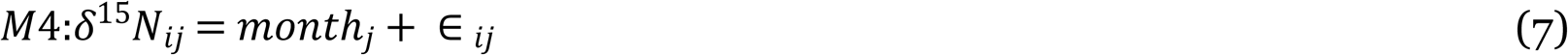

All models were fitted and compared with each other using the corrected Akaike’s Information Criterion (AICc) [48]. All analyses were performed in R 3.1.3 [49].

All applicable international, national, and institutional guidelines for the care and use of animals were followed and all sampling permits and approvals acquired as part of project 839.08.241 of the National Ocean and Coastal Research Programme (ZKO) of the Netherlands Organization for Scientific Research (NWO), for which permit NIOZ 10.03 was given for use of animals by the Animal Experiment Commission of the Royal Netherlands Academy of Sciences (KNAW).

## Results

Mnemiopsis leidyi was not observed in February, and was first observed on March 24, and was observed throughout the rest of the year, except for two weeks in June. Densities remained below 1 ind m-3 until August, when densities started to increase exponentially. Maximum densities were reached in August (mean 63.5 ± 31.0 ind m-3) and in October and November densities had decreased an order of magnitude again (Fig 1). Additional information on *M. leidyi* presence, as well as densities of other gelatinous plankton in 2011 and 2009, 2010 and 2012 can be found elsewhere [38].

**Fig 1.**
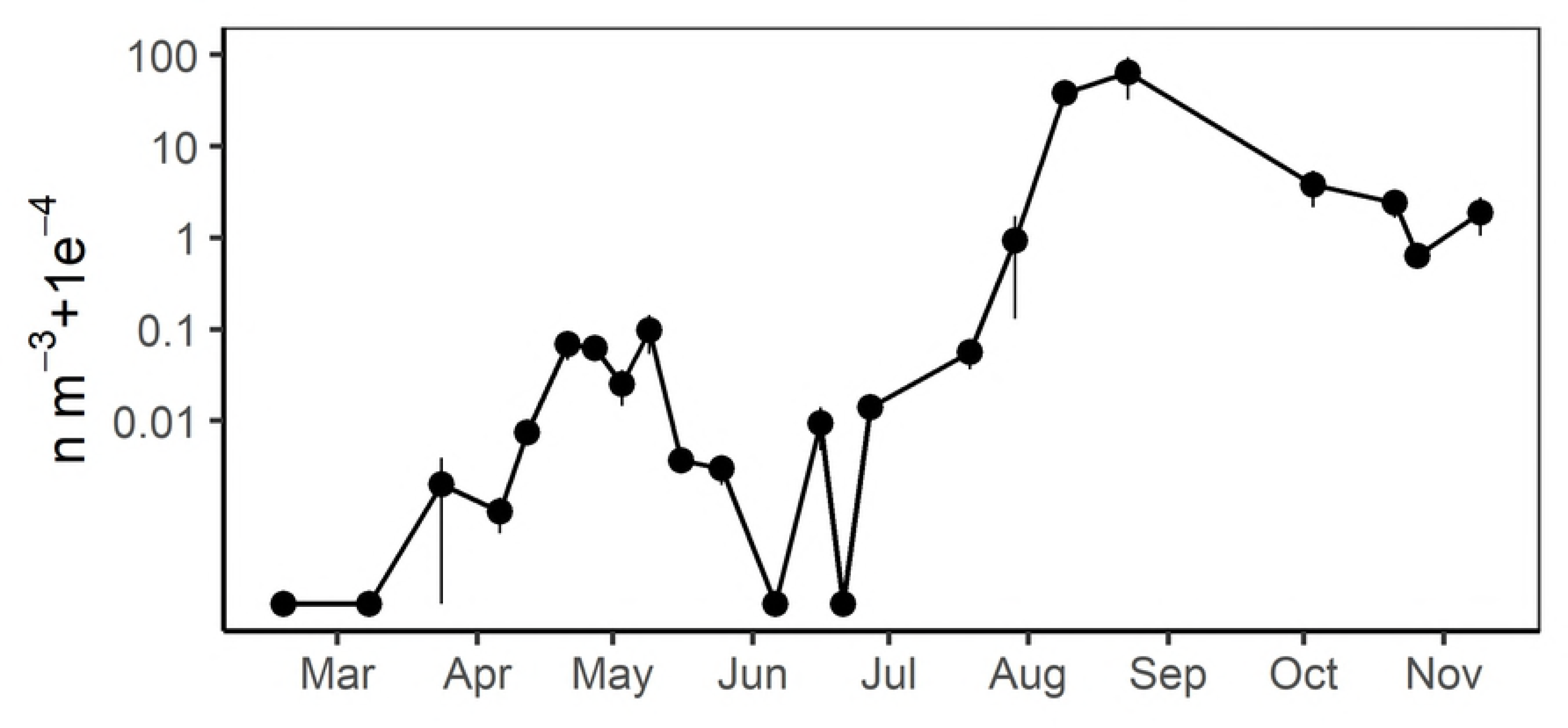
Seasonal pattern of *Mnemiopsis leidyi* density in the western Wadden Sea in 2011. Weekly mean and Standard Error of *Mnemiopsis leidyi* densities (n m^-3^ + 1e^-4^) in 2 mm mesh size gelatinous zooplankton nets in the western Wadden Sea.

*δ*^15^N and *δ*^13^C ratios were obtained for 683 individuals of 36 different species on 26 different sampling days. Most species caught were fish (19 species). 13 species of gelatinous zooplankton were caught: four ctenophores, five scyphomedusae and four hydromedusae. Two amphipods and two mysid species were also sampled (Table 3). Most bulk plankton samples did not analyse properly on the IRMS because the amount of C and N in the samples was too low, and data on the two plankton fractions could only be obtained for August. The baseline *δ*^15^N values from long-lived filter-feeding primary consumers, the bivalve *Mytilus edulis* were collected in March and July 2014 in the Mokbaai (53.0041 N, 4.7711E) and had a mean *δ*^15^N value of 10.8 ‰ and a SE of 0.1 ‰ (n=15). Mean *δ*^15^N values of *M. edulis* did not differ between March and July (one-way ANOVA, *F*_(1,13)_=1.503, *p*=0.242) and were similar to mean *δ*^15^N values found for the >80 µm plankton fraction (one-way ANOVA, *F*_(1,17)_=1.133, *p*=0.302). *δ*^15^N values of filter feeding *Ensis leei* bivalves collected near the main freshwater input source for the area (mean *δ*^15^N 9.6 ‰, SE 0.4 ‰, n = 3) were similar to *δ*^15^N values of bivalves collected near the entrance of the Wadden Sea from the North Sea (mean *δ*^15^N 9.7 ‰, SE 0.2 ‰, n = 8).

**Table 3.** Sizes and means and standard error of δ^13^C, δ^15^N and C:N ratio, per species over the whole sampled period in 2011

| Species | n | size mm | | $\delta^{13}\text{C}$ ‰ | | $\delta^{15}\text{N}$ ‰ | | C:N | |
| --- | --- | --- | --- | --- | --- | --- | --- | --- | --- |
|  |  | min | max | mean | SE | mean | SE | mean | SE |
| *Aequorea vitrina* | 6 | 35 | 81 | -16.6 | 0.8 | 15.4 | 0.5 | 3.5 | 0.1 |
| *Agonus cataphractus* | 1 | 45 | 45 | -17.5 |  | 17.2 |  | 3.3 |  |
| *Alloteuthis subulata* | 5 | 47 | 88 | -18 | 0.3 | 14.7 | 0.2 | 3.7 | 0 |
| *Ammodytes tobianus* | 18 | 44 | 182 | -18.3 | 0.2 | 15.1 | 0.2 | 3.4 | 0.1 |
| *Aphia minuta* | 17 | 25 | 57 | -18.3 | 0.2 | 15.2 | 0.3 | 3.5 | 0 |
| *Aurelia aurita* | 38 | 23 | 251 | -18.5 | 0.2 | 13.5 | 0.2 | 3.4 | 0.1 |
| *Beroe cucumis* | 26 | 15 | 70 | -18.5 | 0.3 | 14.1 | 0.3 | 3.9 | 0 |
| *Beroe gracilis* | 25 | 12 | 34 | -19.4 | 0.5 | 14.5 | 0.2 | 3.8 | 0.1 |
| *Chrysaora hysoscella* | 21 | 15 | 240 | -18.9 | 0.3 | 14.2 | 0.3 | 3.6 | 0.1 |
| *Ciliata mustela* | 5 | 24 | 38 | -19.3 | 0.3 | 16.1 | 0.2 | 3.4 | 0 |
| *Clupea harengus* | 30 | 23 | 139 | -19.1 | 0.2 | 15.3 | 0.2 | 3.4 | 0.1 |
| *Cosmetira pilosella* | 4 | 10 | 16 | -19.7 | 0.1 | 12.5 | 0.2 | 4 | 0.1 |
| *Cyanea capillata* | 6 | 44 | 75 | -18.4 | 0.3 | 13.1 | 0.6 | 3.6 | 0.1 |
| *Cyanea lamarckii* | 18 | 29 | 157 | -17.4 | 0.3 | 13.7 | 0.3 | 3.4 | 0 |
| *Engraulis encrasicolus* | 5 | 33 | 49 | -18.9 | 0.2 | 15.2 | 0.2 | 3.2 | 0 |
| *Gasterosteus aculeatus* | 24 | 22 | 65 | -22.7 | 0.7 | 16.9 | 0.4 | 3.7 | 0.1 |
| *Gastrosaccus spinifer* | 5 | 13 | 15 | -17.7 | 0.3 | 13 | 0.6 | 3.6 | 0 |
| *Hyperia galba* | 25 |  |  | -18.1 | 0.2 | 14.4 | 0.3 | 4 | 0.1 |
| *Hyperoplus lanceolatus* | 8 | 51 | 149 | -19.6 | 0.2 | 14.5 | 0.2 | 3.3 | 0 |
| *Idotea linearis* | 19 | 13 | 24 | -17 | 0.2 | 13.1 | 0.1 | 4.6 | 0.1 |
| *Liparis liparis* | 5 | 43 | 67 | -17.5 | 0.1 | 15.9 | 0.2 | 3.3 | 0 |
| *Loligo vulgaris* | 4 | 15 | 22 | -18.2 | 0.3 | 17.8 | 0.5 | 3.6 | 0 |
| *Merlangius merlangus* | 5 | 46 | 120 | -18.1 | 0.3 | 17.1 | 0.3 | 3.2 | 0 |
| *Mnemiopsis leidyi* | 74 | 6 | 72 | -19 | 0.1 | 14.5 | 0.2 | 4 | 0 |
| *Nemopsis bachei* | 19 | 7 | 13 | -19.3 | 0.1 | 13.8 | 0.2 | 3.9 | 0 |
| *Noctiluca scintillans* | 1 |  |  | -20.3 |  | 10.3 |  | 6.8 |  |
| *Osmerus eperlanus* | 15 | 52 | 142 | -17.9 | 0.4 | 17.2 | 0.2 | 3.2 | 0 |
| *Pleurobrachia pileus* | 75 | 7 | 30 | -18.3 | 0.1 | 14.7 | 0.1 | 3.6 | 0 |
| *Pleuronectes platessa* | 5 | 9 | 14 | -23.1 | 0.4 | 15.2 | 0.9 | 4 | 0.1 |
| *Pomatoschistus lozanoi* | 13 | 19 | 58 | -18.2 | 0.3 | 15.9 | 0.2 | 3.3 | 0 |
| *Pomatoschistus minutus* | 15 | 24 | 82 | -18.5 | 0.4 | 16.3 | 0.1 | 3.2 | 0 |
| *Praunus flexuosus* | 6 | 18 | 26 | -16.1 | 0.1 | 14.5 | 0.3 | 3.3 | 0 |
| *Rhizostoma octopus* | 19 | 15 | 130 | -19.8 | 0.1 | 13.6 | 0.2 | 3.5 | 0.1 |
| *Sarsia tubulosa* | 9 | 10 | 11 | -20.2 | 0.1 | 12.3 | 0.2 | 3.9 | 0 |
| *Sprattus sprattus* | 25 | 39 | 132 | -18.6 | 0.1 | 14.3 | 0.2 | 3.4 | 0.1 |
| *Syngnathus rostellatus* | 19 | 30 | 139 | -18.7 | 0.1 | 15.5 | 0.3 | 3.5 | 0.1 |
| *Trachurus trachurus* | 10 | 11 | 50 | -19.8 | 0.1 | 16.1 | 0.3 | 3.3 | 0 |
| *Trisopterus luscus* | 2 | 80 | 135 | -17.4 | 0.5 | 17.6 | 0.2 | 3.3 | 0 |
| plankton fraction (> 80 µm) | 5 |  |  | -16.8 | 0.4 | 10.6 | 0.5 | 7.1 | 0.4 |
| plankton fraction (< 80 µm) | 9 |  |  | -14.6 | 1.4 | 8.2 | 0.4 | 8.6 | 0.6 |

The average *δ*^15^N of all species is shown in Table 3 and ranked in Fig 2. The squid *Loligo vulgaris* had the highest mean *δ*^15^N ratio at 17.8 ± 0.5 ‰ followed by several fish species such as the bib *Trisopterus luscus*, the hooknose *Agonus cataphractus*, the whiting *Merlangius merlangius* and the smelt *Osmerus eperlanus*. The gelatinous animal with the highest mean *δ*^15^N ratio was *Aequorea vitrina*, which had the 12^*th*^ highest value overall; 15.4 ± 0.5 ‰. The scyphomedusae species with the highest mean *δ*^15^N ratio was *Chrysaora hysoscella* at 14.2± 0.3 ‰. The other scyphomedusae species had a surprisingly low mean *δ*^15^N ratio of less than 14 ‰.

**Fig 2.**
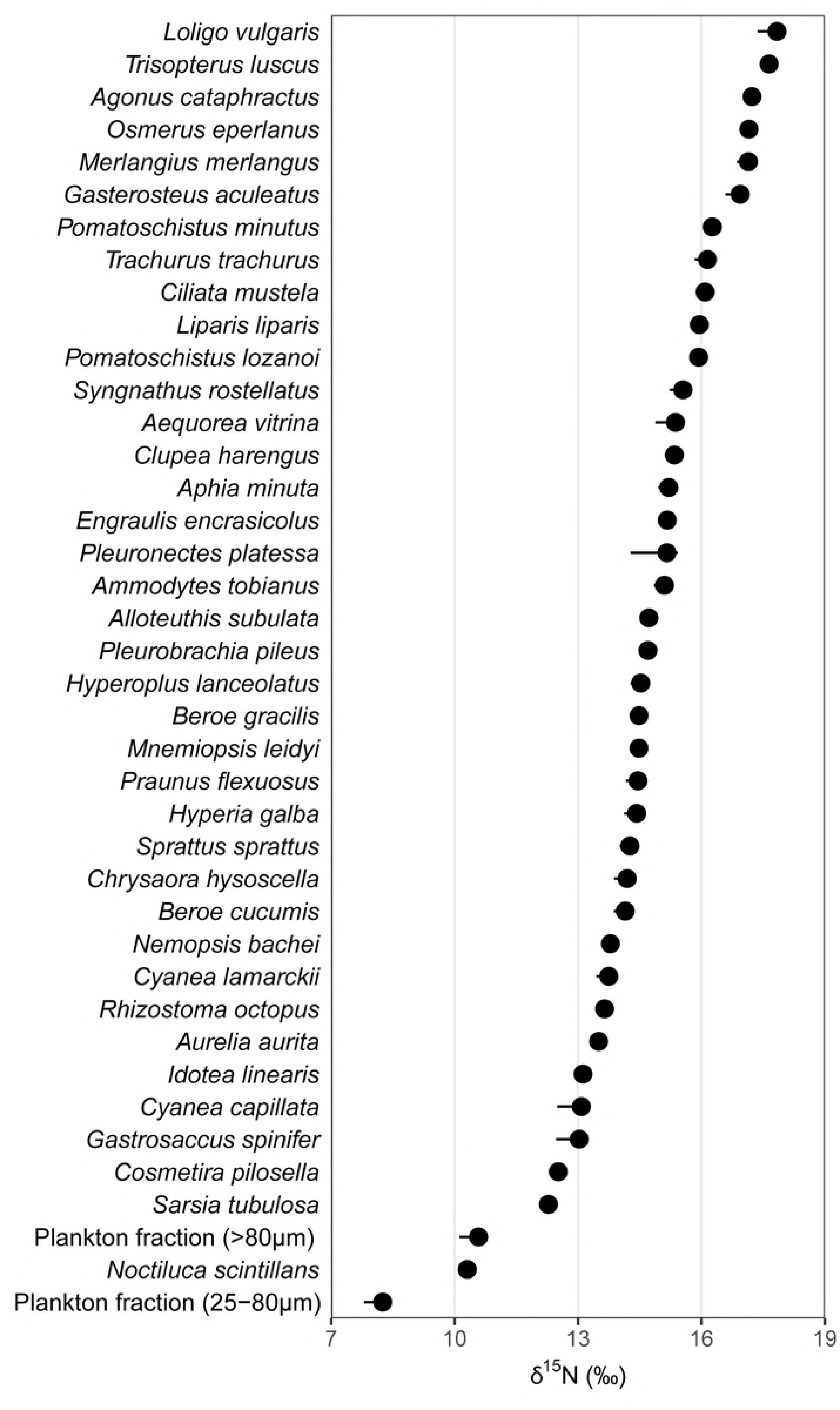
Average nitrogen stable isotope ratios of sampled taxa in 2011. Mean and Standard Error of δ^15^N per species in 2011, ordered highest to lowest

Average of *Mnemiopsis leidyi* was 14.5 ± 0.2 ‰, close to that of *Pleurobrachia pileus* (14.7 ± 0.1 ‰), *Beroe gracilis* (14.5 ± 0.2 ‰) and *Beroe cucumis* (14.1 ± 0.3 ‰). Gelatinous zooplankton species with the lowest *δ*^15^N ratios were the hydromedusae *Sarsia tubulosa* and *Cosmetira pilosella*. The lowest values were found for the <80 µm plankton fraction (8.2 ± 0.4 ‰) which was close to the minimum value of 1 for autotrophs. The mixotrophic dinoflagellate *Noctiluca scintillans* had a *δ*^15^N ratio of 10.3 ‰.

### Isotopic niches

The shape and surface area of the Standard Ellipse Area corrected for small sample sizes (SEAc) gives information on niche width of consumers, whereby the SEAc of a generalist is larger than that of a specialist feeder. Isotopic niches were both estimated for species and for higher taxonomic ranks.

### Isotopic niches of higher taxonomic ranks

Species were grouped according to taxonomical groups fish, cephalopods, crustaceans, ctenophores, hydromedusae and Scyphomedusae, and for each group the isotopic niche in the form of the Standard Ellipse Area corrected for small sample size (SEA_c_) was estimated (Table 2 and Fig 3). The isotopic niche of fish overlapped most with that of the cephalopods and ctenophores, and least with that of scyphozoa and crustacea. Fish had the largest isotopic niche, followed by the ctenophores, hydromedusae, scyphomedusae, crustacea and cephalopods.

**Table 2.**
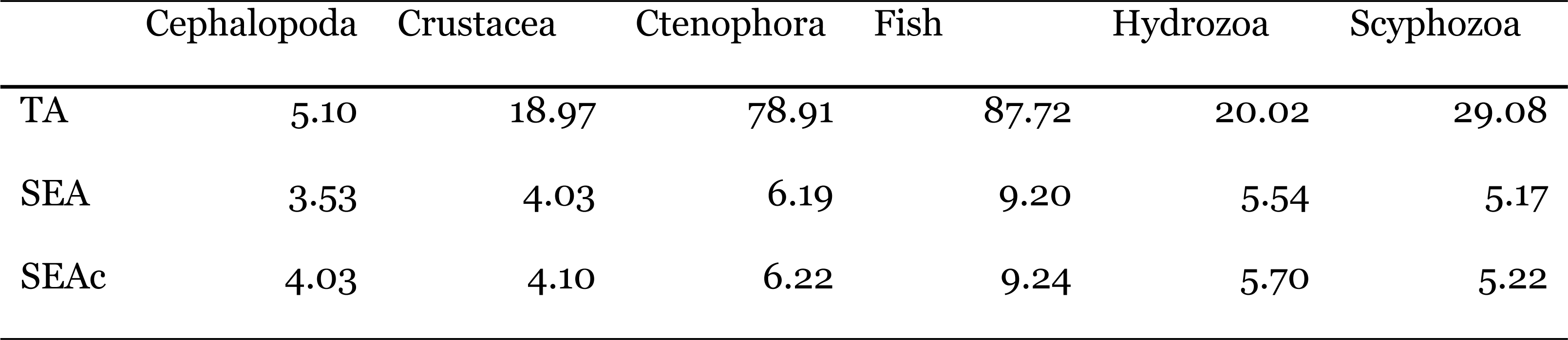
Total area of the convex hull encompassing the data points (TA), Standard Ellipse Area (SEA) and Standard Ellipse Area corrected for small sample sizes (SEA_c,_), all expressed as ‰^2^).

**Fig 3.**
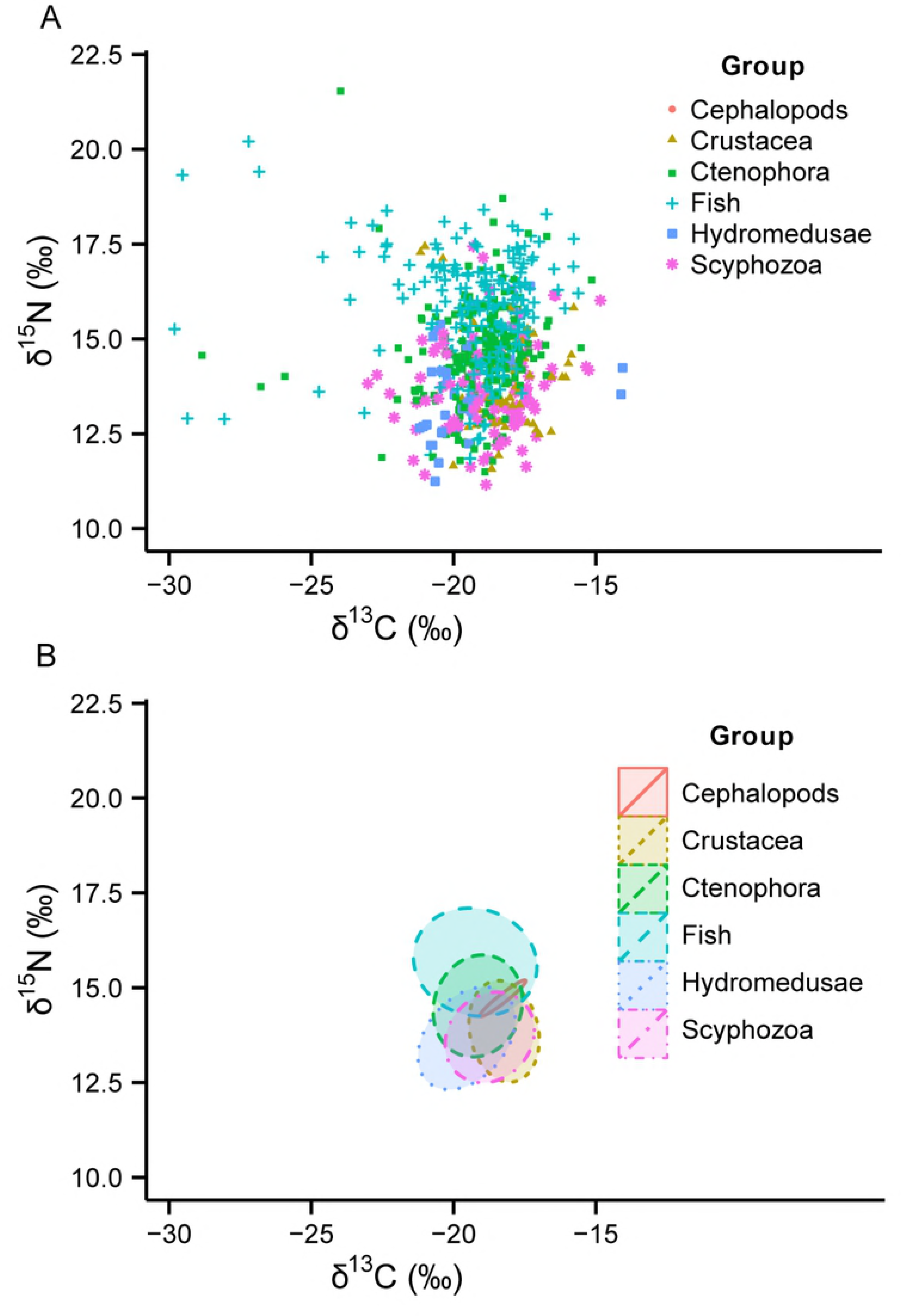
Biplots of carbon and nitrogen stable isotope ratios and of samples in each taxonomical group and their estimated isotopic niches. (A) δ^15^N and δ^13^C values of consumers by taxonomical group and (B) niche of the total group estimated as the Standard Ellipse Area corrected for small sample size (SEA_c_)

### Isotopic niches of species

SEA_*c*_ was very variable between species as well as within species. The SEA_*c*_ of several pelagic consumers overlapped with that of *M. leidyi* in one or more months (Table 4). The decrease in *δ*^15^N of *M. leidyi* over time and with decreasing length meant that isotopic niche overlap of *M. leidyi* with other species was not constant over the whole studied period but differed each month. SEA_*c*_ overlap was highest in the spring months, when the SEA_*c*_ of *M. leidyi* overlapped with that of several pelagic fish species, ctenophores, scyphomedusae, hydromedusae and cephalopods.

**Table 4.** Standard Ellipse Area corrected for small sample sizes (SEA_c_, as δ units^2^) with percent overlap of the SEA_c_ of consumers with that of *Mnemiopsis leidyi* in parentheses, for each month and for all months combined(overall)

| Species | Month |  |  |  |  |  | overall |
| --- | --- | --- | --- | --- | --- | --- | --- |
|  | March | April | May | June | July | August |  |
| *Alloteuthis subulata* |  |  | 0.43 (47) |  |  |  | 0.43 (51) |
| *Ammodytes tobianus* | 0.67 (0) | 2.22 (9) | 0.44 (2) |  |  |  | 2.44 (62) |
| *Aphia minuta* | 3.07 (38) | 2.68 (22) | 0.76 (42) |  |  |  | 2.72 (59) |
| *Aurelia aurita* |  | 3.71 (0) | 1.45 (0) | 1.96 (6) |  |  | 3.68 (38) |
| *Beroe cucumis* |  |  | 0.52 (0) | 1.06 (12) |  | 0.37 (0) | 2.19 (51) |
| *Beroe gracilis* |  |  | 1.02 (29) | 0.87 (68) |  | 2.34 (0) | 4.48 (66) |
| *Chrysaora hysoscella* |  |  |  | 3.13 (70) |  | 1.97 (0) | 3.11 (94) |
| *Ciliata mustela* |  |  |  | 0.69 (100) |  |  | 0.69 (41) |
| *Clupea harengus* | 3 (58) | 4.29 (10) | 0.24 (4) | 3.46 (73) |  |  | 4.33 (70) |
| *Cyanea lamarckii* |  |  | 1.47 (0) | 3.17 (0) |  |  | 2.57 (5.8) |
| *Engraulis encrasicolus* |  |  |  |  |  | 0.8 (0) | 0.80 (99) |
| *Gasterosteus aculeatus* | 1.99 (0) | 3.43 (0) | 1.52 (0) | 27.89 (0) |  |  | 17.39 (0) |
| *Gastrosaccus spinifer* |  |  |  |  |  | 3.13 (0) | 3.13 (1) |
| *Hyperia galba* |  |  | 2.06 (0) | 1.7 (55) |  |  | 2.55 (35) |
| *Hyperoplus lanceolatus* |  | 0.57 (5) |  |  |  |  | 0.57 (100) |
| *Idotea linearis* |  | 1.42 (0) | 0.56 (0) | 0.09 (0) |  |  | 0.94 (0) |
| *Liparis liparis* |  |  |  | 0.3 (0) |  |  | 0.30 (0) |
| *Merlangius merlangus* |  |  |  | 1.05 (0) |  |  | 1.05 (0) |
| *Mnemiopsis leidyi* | 5.52 | 1.02 | 0.5 | 5.81 | 1.38 | 0.39 | 5.21 |
| *Nemopsis bachei* |  |  |  | 2.00 (57) | 1.49 (29) | 0.25 (0) | 1.69 (98) |
| *Osmerus eperlanus* |  |  | 0.25 (0) |  |  |  | 0.25 (0) |
| *Pleurobrachia pileus* | 4.34 (29) | 2.45 (27) | 0.99 (0) | 1.61 (24) | 3.68 (0) | 0.48 (0) | 4.09 (64) |
| *Pleuronectes platessa* | 6.65 (0) |  |  |  |  |  | 6.65 (0) |
| *Pomatoschistus lozanoi* |  |  | 0.20 (0) |  |  |  | 0.20 (85) |
| *Pomatoschistus minutus* |  |  | 3.63 (0) |  |  |  | 3.63 (4) |
| *Praunus flexuosus* |  |  |  |  |  | 0.54 (0) | 0.54 (0) |
| *Rhizostoma octopus* |  |  |  | 1.2 (0) |  |  | 1.20 (30) |
| *Sarsia tubulosa* | 0.49 (0) |  |  |  |  |  | 0.49 (0) |
| *Sprattus sprattus* |  | 0.93 (0) | 1.2 (16) | 1.83 (41) |  | 0.29 (0) | 1.75 (99) |
| *Syngnathus rostellatus* |  |  | 1.81 (0) | 0.9 (2) |  | 0.33 (0) | 2.62 (68) |
| *Trachurus trachurus* |  |  |  | 0.62 (94) |  | 0.63 (0) | 1.19 (25) |
| Overlapping species | 3/7 | 4/9 | 6/17 | 12/18 | 1/2 | 0/11 | 24/30 |

Fish species which had high percentages of SEA_*c*_ overlap with *M. leidyi* were the glass goby *Aphia minuta*, the five-beard rockling *Ciliata mustela*, the herring *Clupea harengus*, the sand eel *Hyperoplus lanceolatus*, the lesser pipefish *Syngnathus rostellatus*, the sprat *Sprattus sprattus* and the horse mackerel *Trachurus trachurus*. Gelatinous zooplankton species with overlapping SEA_*c*_ with *M. leidyi* were mainly *Beroe gracilis*, *Chrysaora hysoscella*, *Pleurobrachia pileus* and *Nemopsis bachei*. In August no overlap in SEA_*c*_ of any species with that of *M. leidyi* was observed.

### Clusters

All ctenophores and scyphomedusa clustered together with most pelagic fish species of intermediate trophic position (Fig 4) including herring *Clupea harengus* and sprat *Sprattus sprattus* in cluster A. This cluster also included the squid *Alloteuthis subulata*, the parasitic amphipod *Hyperia galba*, an isopod(*Idotea linearis*) and amysid(*Gastrosaccus spinifer*). *Mnemiopsis leidyi* was grouped together in a subcluster with the ctenophore predator *Beroe gracilis* and the sand eel *Hyperoplus lanceolatus*. Cluster B contained the hydromedusa *Aequorea vitrina* and the mysid *Praunus flexuosus*. Cluster C contained several fish species that occupied the highest trophic positions, as well as the squid *Loligo vulgaris*. Cluster D contains the dinoflagellate *Noctiluca scintillans* and interestingly also two hydroid species, *Sarsia tubulosa* and *Cosmetira pilosella*. Pelagic larvae of plaice *Pleuronectes platessa* were grouped together with the anadromous three-spined stickleback *Gasterosteus acuelatus* in cluster E. Branch F and G consisted solely the different bulk zooplankton fractions.

**Fig 4.**
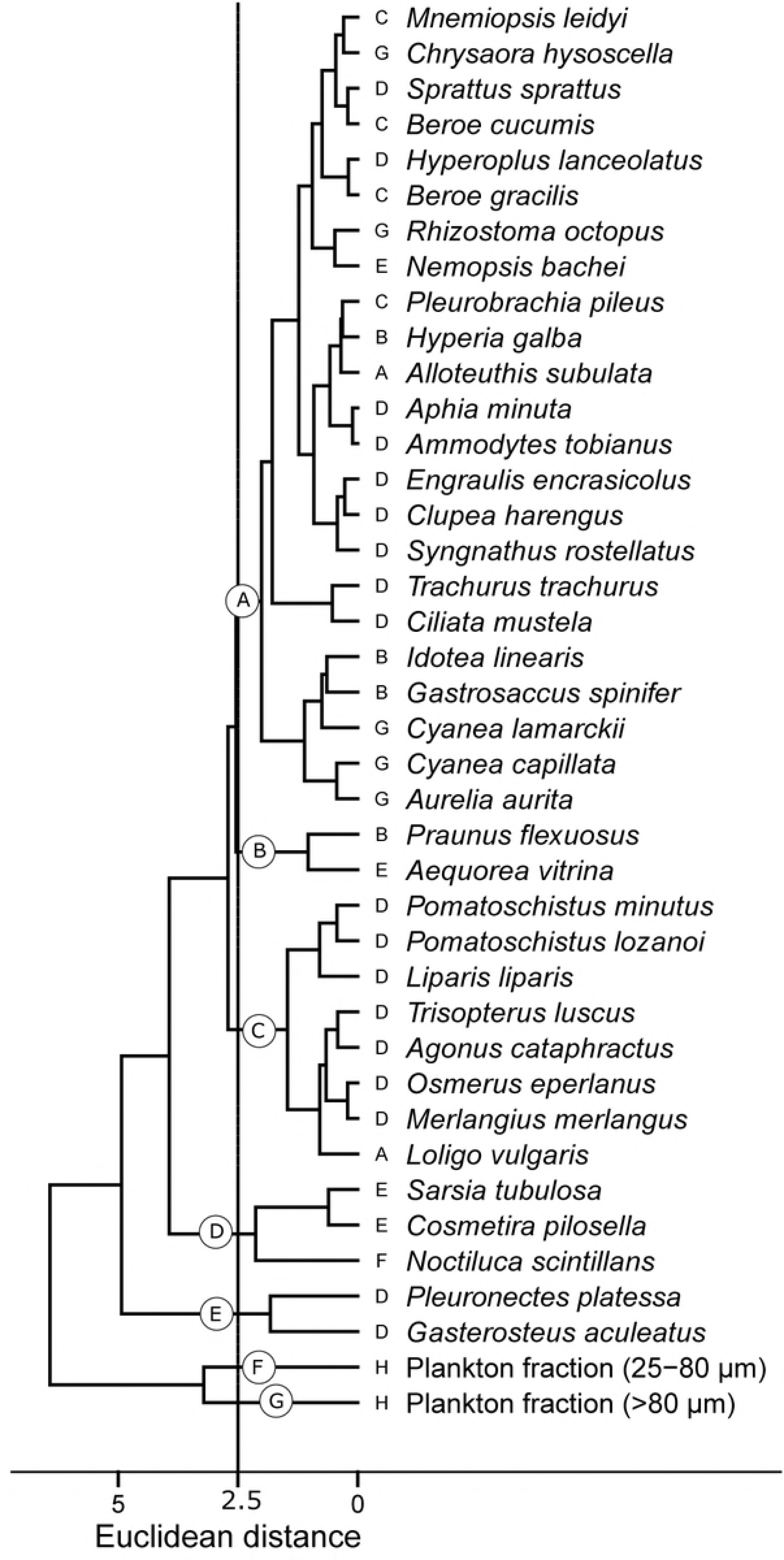
Hierarchical cluster analysis of western Wadden Sea pelagic consumers in 2011. Euclidean metric distances calculated with the group-average method based on mean *δ*^15^N and δ13C values for each species. Capital letters in circles indicate the main clusters at 2.5 Euclidean metric distance. Taxonomical groups indicated by capital letters: (A) Cephalopods, (B) Crustacea, (C) Ctenophora, (D) Fish, (E) Hydromedusae, (F) Phytoplankton, (G) Scyphozoa, (H) Zooplankton.

The cluster analysis was also performed for each month separately. As not all species were caught in each month and stable isotope values varied within species, the cluster tree topology was very different in each month. In March–June *M. leidyi* clustered together with mainly zooplanktivorous fish such as *C. harengus*. In July, when fewer species could be sampled, no different clusters could be discerned at 2.5 Euclidean Metric Distance. In August *M. leidyi* did not cluster together with *C. harengus* anymore and occupied a sub-cluster with *Cosmetira pilosella* and *B. gracilis*.

### Seasonal patterns in Stable Isotope ratios

*δ*^15^N and *δ*^13^C of most gelatinous zooplankton species varied considerably over the studied period (Fig 5). For ctenophores *δ*^15^N was almost constant over the first three months, after which there was an increase for both *Beroe cucumis* and *Pleurobrachia pileus* in June, followed by a decrease in all species in July and August. *B. cucumis* appeared in May and had a 2 ‰ lower *δ*^15^N ratio than the other species in this month. Also for *B. gracilis δ*^15^N ratios were lower than those of its potential prey species *M. leidyi* and *P. pileus* in several months. During the bloom of *M. leidyi* in July and August, when many small ctenophores were present, *δ*^15^N of both *Beroe* species decreased at the same time as the *δ*^15^N of *M. leidyi* decreased, while the *δ*^15^N of *P. pileus* remained constant and ended up being higher than those of the predatory *Beroe* species in August.

**Fig 5.**
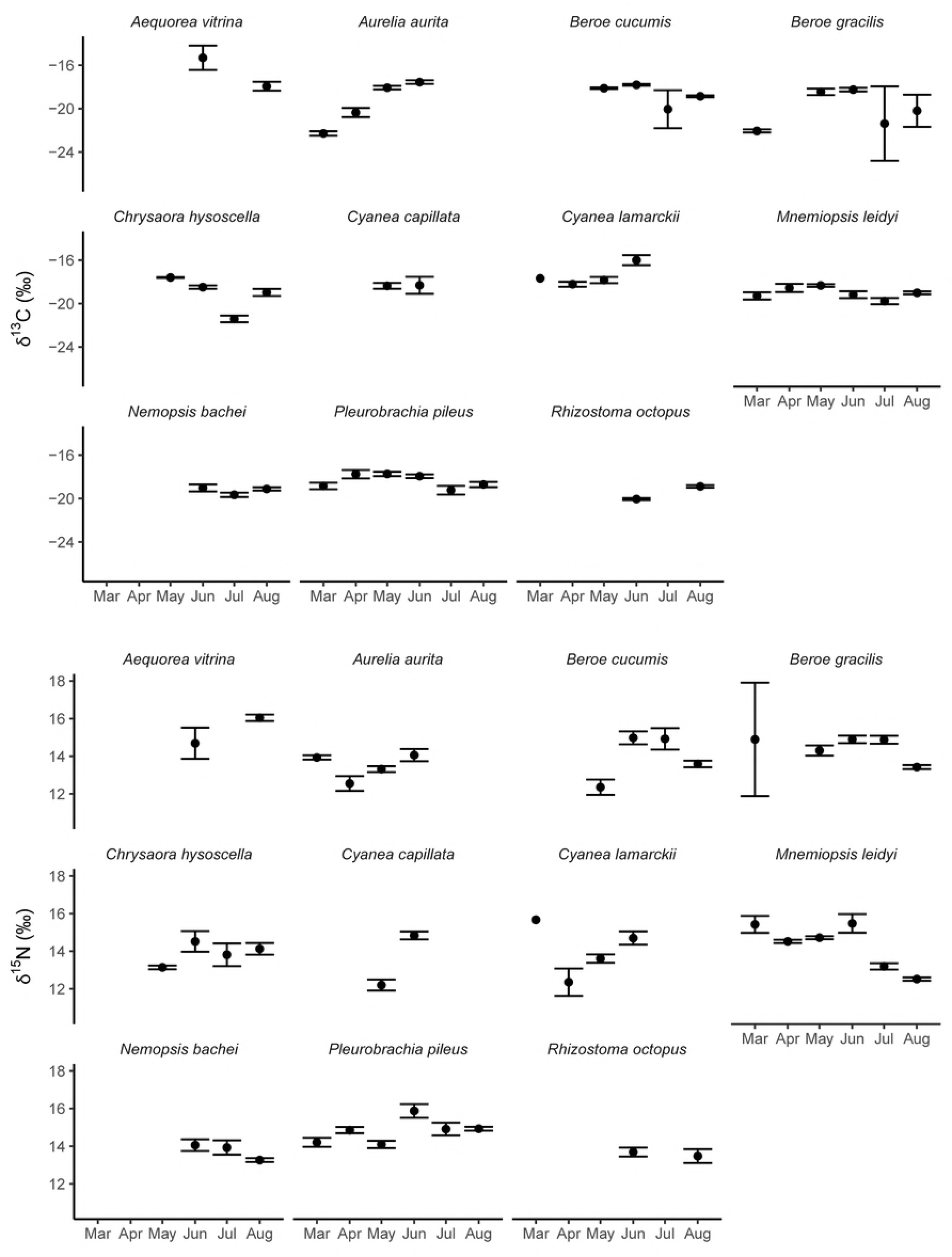
Mean monthly carbon and nitrogen stable isotope ratios in western Wadden Sea gelatinous zooplankton in 2011. Average (+/- SE) δ^13^C (top) and δ^15^N (bottom) per month sampled for all gelatinous species in 2011, excluding *Sarsia tubulosa* and *Cosmetira pilosella* which were only observed once.

*Sarsia tubulosa* was only caught in March. *δ*^15^N of *Aequorea vitrina* increased while that of *Nemopsis bachei* decreased from June to August. *Cosmetira pilosella* was only caught in August.

In March and April only *Cyanea lamarckii* and *Aurelia aurita* were present, which had similar *δ*^15^N ratios. The increase in *δ*^15^N ratios as observed in the ctenophores in June was also observed in all scyphozoa species except *Rhizostoma octopus* which appeared in June. *R. octopus* and *Chrysaora hysoscella* where the only species present in July and August, and they had similar *δ*^15^N ratios in this period.

Patterns in *δ*^13^C were less variable in ctenophores. In March *B. gracilis* was depleted compared to its prey. All species of ctenophores showed a similar seasonal pattern, with depletion occurring between June and July, followed by a slight enrichment in August. The hydromedusae *A. vitrina* was the most enriched species and *Cosmetira pilosella* the most depleted one. *δ*^13^C values of all species occurring in June - August were different. *δ*^13^C-of both *A. aurita*-and *C. lamarckii* increased from March May. From June July both species present showed depletion in *δ*^13^C, the highest in *R. octopus*, followed by enrichment as was also observed in the ctenophores.

### Isotopic position of *Mnemiopsis leidyi*

For *M. leidyi* the relationship between ctenophore oral-aboral length and *δ*^15^N for different months was best described by the model without interaction between month and size (M2, r^2^ = 0.50, *F*_6,67_ = 11.35, *p* < 0.0001) with a different intercept but similar slope for the relationship between *δ*^15^N and ctenophore length in each month. This model had the lowest AICc of the four tested models. Residuals were normally distributed but outliers were present, which all had Cook’s Distance values of less than 0.5. There was a significant (p <0.05) relationship between *δ*^15^N and ctenophore length with a different intercept for each month (Fig 6). *δ*^15^N increased with increasing *M. leidyi* length.

**Fig 6.**
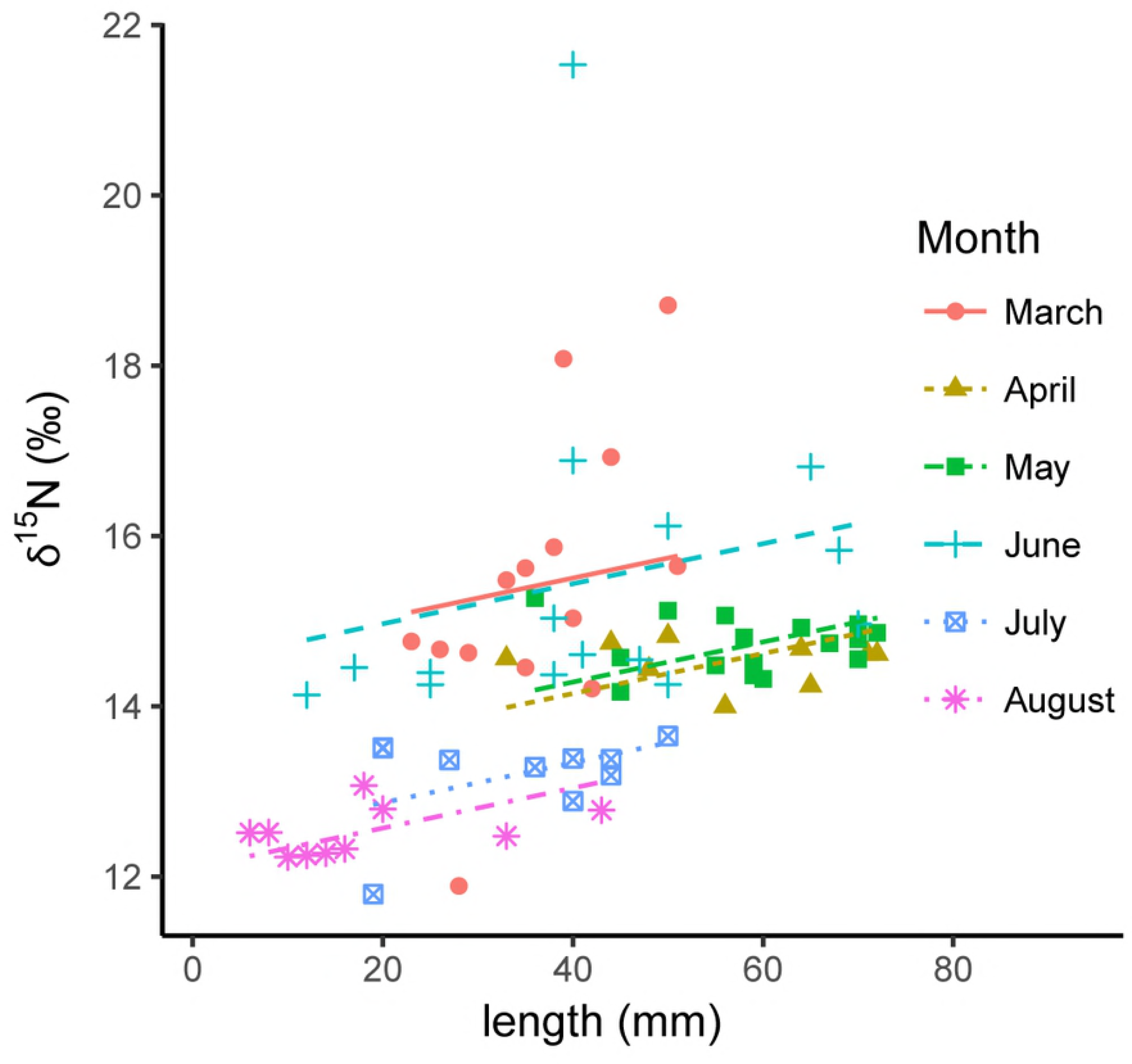
Observed and expected relationship between *Mnemiopsis leidyi* nitrogen stable isotope ratio and ctenophore length. Model results for linear model M1 for *Mnemiopsis leidyi* showing the relationship between δ15N and length in different months in 2011, together with observed values.

## Discussion

### Baseline stable isotope ratios

The western Dutch Wadden Sea is under the influence of freshwater and nutrient input from various sources [50] as well as coastal water input from the nearby North Sea. The average residence time of the water in the Marsdiep tidal basin is ca. 8.5 days [51], but residence time and flushing rate is highly dependent on wind conditions [52]. Consequently, species caught in the western Wadden Sea might have spent part of their life feeding in the nearby North Sea, where mesozooplankton *δ*^15^N and *δ*^13^C can be lower [27,53,54]. For these species our baseline of filter feeding bivalves from within the area might not be appropriate. *Mnemiopsis leidyi* and *Clupea harengus* carbon and nitrogen stable isotope ratios were similar to that found in the eastern Wadden Sea [22], except for some *M. leidyi* individuals larger than 40 mm, which had higher *δ*^15^N values. Stable isotope ratios of filter feeding bivalves used as baseline were also similar between eastern and western Wadden Sea.

The large inputs of anthropogenic nitrogen that enters the western Wadden Sea via several freshwater sources [50] might also influence the baseline of the system. In Naragansett Bay a clear spatial gradient could be seen in *δ*^15^N enrichment of macroalgae, with *δ*^15^N becoming more depleted further away from the source, and *δ*^15^N of clams collected in the bay being enriched compared to clams collected outside of the bay, suggesting that the clams feed on phytoplankton supported by anthropogenic N [55]. In the Wadden Sea, the *δ*^13^C of consumers feeding on pelagic producers was homogeneous across the whole area [56] and filter feeding *Ensis leei* bivalves collected for this study showed similar *δ*^15^N values both close to the freshwater input source as near the entrance to the North Sea, suggesting that at least within the Wadden Sea the baseline stable isotope ratios are similar. Regardless, variation in baselines between different possible source areas for could imply that variation in *δ*^15^N might not be a result of species feeding on different trophic levels. The addition of a third isotope, sulfur, could help to disentangle sources with similar C and N isotope ratios [57].

### Isotopic niches of gelatinous zooplankton and small pelagic fish in the Wadden Sea

While most of the highest δ15N values were of fish, a part of the isotopic niche of fish overlapped with that of ctenophores, hydromedusae and scyphomedusae. The cluster analyses show that stable isotopes signatures of many fish species are similar to those of gelatinous zooplankton, which means that all these groups have to be included when assessing the grazing pressure on zooplankton and possible competition for food between fish and other species.

For many species, stable isotope ratios varied over time, which resulted in both the Standard Ellipse Area comparisons as well as the cluster analyses showing large differences between months, suggesting that diet is not constant over time in the western Wadden Sea. This has been observed in gut content analyses of several fish species in the eastern Wadden Sea as well [22]. This study highlights that isotopic niches of gelatinous zooplankton, including *M. leidyi* can vary between species but also within species with size and season, as found in other areas as well [58,59].

Interestingly, carbon and nitrogen stable isotope ratios of several species that appeared in late spring t o early summer were quite different from those of species that had already been present for several months. The most striking example of this is *Beroe cucumis* which was depleted in *δ*^15^N compared to the other ctenophore species when it appeared in May, followed by enrichment to similar levels in the next month.

A possible explanation for this is that *Beroe cucumis* appeared in the Wadden Sea by advection from the nearby North Sea, where it had been feeding on its preferred prey, the lobatē ctenophore *Bolinopsis infundibulum* which has been shown to be depleted in *δ*^15^N compared to other species of ctenophores [53]. A similar deviation of stable isotope ratios in *Beroe* was found in the southern North Sea [60]. When *Beroe* entered the Wadden Sea it could have started feeding on the species present there, consequently becoming enriched in *δ*^15^N. Something similar might have occurred for *Cyanea capillata* from May - June as well as this is also a species that is known to be more abundant in the North Sea than close to the coast [61]. The native ctenophore *Beroe gracilis* has been shown to be able to prey successfully on *M. leidyi* in experiments but *M. leidyi* of 20 mm oral-aboral length or larger could only be consumed partially by *B. gracilis* [62]. In this study the *δ*^15^N of *Beroe* species was often equal or lower than that of *M. leidyi* and that of its main prey, *P. pileus*. After high densities of small (<10 mm) sized *M. leidyi* started to dominate the catches the *δ*^15^N of both *Beroe* species decreased suggesting that small *M. leidyi* is their main prey in this period.

### Variation in diet of *Mnemiopsis leidyi*

Adult *Mnemiopsis leidyi* can cruise through the water by rhythmic beating of their ciliary comb rows, and generate a feeding current past their tentillae [63]. This current is difficult to detect by most prey species until they are well between the oral lobes [63]. Prey items that do not have an escape response to the feeding current are retained directly onto the tentillae, while fast-moving prey such as copepods are caught on the inside surface of the lobes [64]. Consequently *M. leidyi* is a generalist, feeding on a wide range of prey from microplankton to copepods and fish larvae [3].

The isotopic niche of *M. leidyi* overlapped with those of several fish species, including the herring *Clupea harengus*. A similar overlap in diet between *C. harengus* and *M. leidyi* was found earlier in the eastern Wadden Sea [21] and the division of the fish species in this study is largely in agreement with a similar cluster analysis of eastern Wadden Sea zooplanktivores [22], with the fish species clustering in cluster C in this study belonging to a guild of mostly *Crangon crangon* eaters, and the species in cluster A belonging to a guild feeding mainly on calanoid copepods. The fish species that showed the most overlap in diet with *M. leidyi* in this study all belonged to cluster A.

Stable isotope ratios of *M. leidyi* also overlapped with those of other gelatinous zooplankton species. These were mainly *Pleurobrachia pileus*, *Chrysaora hysoscella* and *Nemopsis bachei*, all species belonging to the same cluster A.

A striking observation was the lack of isotopic niche overlap of any consumer considered in this study with *M. leidyi* in August at the beginning of its bloom period [17,38] when almost the entire population consisted of larvae and juveniles. The positive relationship between *δ*^15^N and size of *M. leidyi* found in this study suggests that smaller ctenophores have a lower trophic position than larger ones. It has indeed been observed that larval *M. leidyi* are feeding on microplankton [65,66] whereby the relative proportion of microplankton in the diet of *M. leidyi* increases with increasing size, with microplankton contribution to the diet of ctenophores smaller than 10 mm above 80% [67]. Adult ctenophores can decrease competition for their larvae by feeding on microplankton-grazing meso-zooplankton[68]. Additionally, regardless of size, *M. leidyi* has been shown to prey on shellfish larvae in its native habitat [69] and in the eastern Wadden Sea as well [22]. In the western Wadden Sea shellfish larvae were found in M. leidyi stomachs as well, identified using molecular methods as belonging to *Magallana. gigas, Mytilus edulis, Cerastoderma edule, Mya arenaria* and *Ensis leei* [70].

During blooms when the *M. leidyi* population consists mainly of larvae and juveniles, the species might be a competitor rather than a predator of mesozooplankton. The Scyphomedusa *Rhizostoma octopus* feeds mainly on microplankton such as tintinnids [71] and is abundant in late summer and autumn in the western Wadden Sea area [38,72], making it a likely competitor of *M. leidyi*, although in our study it had stable isotope ratios comparable to those of the other Scyphomedusae.

In a related study where mesozooplankton clearance rates of the major gelatinous predators in the western Wadden Sea before and after the introduction of *M. leidyi* were compared, the clearance rates by gelatinous zooplankton in September – November were an order of magnitude higher in 2009 – 2012 with *M. leidyi* present than they were in 1981 – 1983 with *M. leidyi* absent [38]. The period of high diet overlap with other consumers, March – June,was also the period in which *M. leidyi* was least abundant and its estimated zooplankton clearance rates were low [38] suggesting that in 2011 *M. leidyi* was not a significant competitor for food for other mesozooplanktivores in the western Wadden Sea, as has been observed in the eastern Wadden Sea in 2010 [22].

## Acknowledgements

Thanks are due to the crew of the research vessels ‘Stern’ and ‘Navicula’ for their assistance during sampling, to Daphne Rekers for her assistance with the stable isotope analyses, and to the students that assisted with sampling.

